# GFI1 tethers the NuRD complex to open and transcriptionally active chromatin in myeloid progenitors

**DOI:** 10.1101/2021.05.31.446398

**Authors:** Anne Helness, Jennifer Fraszczak, Charles Joly-Beauparlant, Halil Bagci, Christian Trahan, Kaifee Arman, Peiman Shooshtarizadeh, Riyan Chen, Marina Ayoub, Jean-François Côté, Marlene Oeffinger, Arnaud Droit, Tarik Möröy

## Abstract

GFI1 is a SNAG-domain, DNA binding transcriptional repressor which controls myeloid differentiation, in particular the formation of neutrophils. Here we show that GFI1 interacts with the chromodomain helicase CHD4 and other components of the “Nucleosome remodeling and deacetylase” (NuRD) complex. In granulo-monocytic precursors, GFI1, CHD4 or GFI1/CHD4 complexes occupy sites of open chromatin enriched for histone marks associated with active transcription suggesting that GFI1 recruits the NuRD complex to target genes that are regulated by active or bivalent promoters and active enhancers. Our data also show that GFI1 and GFI1/CHD4 complexes occupy promoters of different sets of genes that are either enriched for IRF1 or SPI-1 consensus sites, respectively. During neutrophil differentiation, overall chromatin closure and depletion of H3K4me2 occurs at different degrees depending on whether GFI1, CHD4 or both are present, indicating that GFI1 affects the chromatin remodeling activity of the NuRD complex. Moreover, GFI1/CHD4 complexes regulate chromatin openness and histone modifications differentially to enable regulation of target genes affecting the signaling pathways of the immune response or nucleosome organization or cellular metabolic processes.

## INTRODUCTION

The DNA binding zinc finger proteins GFI1 and GFI1B act as transcriptional repressors by recruiting the LSD1/CoREST and HDACs to sites of specific target genes that harbor a GFI/GFI1B consensus DNA binding motif (Saleque *et al*, 2007) (for a review see (Duan & Horwitz, 2003a; Fraszczak & Moroy, 2017; Moroy *et al*, 2015; van Bergen & van der Reijden, 2019)). GFI1 is critical for differentiation of myeloid cells into neutrophils, which is highlighted for example by GFI1 deficient mice that entirely lack neutrophil granulocytes and, as a consequence, show major defects in their innate immune response (Hock *et al*, 2003; Karsunky *et al*, 2002). GFI1 and its shorter paralog GFI1B share a 20 aa N-terminal SNAG domain that shows sequence similarity to the N-terminal tail of histone H3 (Lin *et al*, 2010). It has been suggested that the GFI1/B SNAG domains and the H3 N-terminus can compete for binding to the same pocket in the LSD1 protein (Lin *et al*., 2010). Transcriptional repression is achieved by the enzymatic action of LSD1 and HDACs leading to the demethylation of H3K4 and the deacetylation of H3K9 (Duan & Horwitz, 2003b; Maiques-Diaz & Somervaille, 2016; Marabelli *et al*, 2016; Saleque *et al*., 2007). The general applicability of this model has been challenged by recent observations indicating that H3K4 methylation states do not change in cells upon treatment with an LSD1 inhibitor that not only blocks its enzymatic activity but also leads to the eviction of GFI1 and LSD1 from promoter sites (Maiques-Diaz *et al*, 2018a; Maiques-Diaz *et al*, 2018b). Moreover, this report finds that LSD1’s demethylase function is not critical for GFI1 function but rather suggests that LSD1’s physical interaction with GFI1’s SNAG-domain is crucial and that LSD1 rather serves as a scaffold for other histone modifying enzymes such as HDACs (Maiques-Diaz *et al*., 2018a; Maiques-Diaz *et al*., 2018b).

To understand the precise molecular function of GFI1 as a transcriptional regulator, it is necessary to identify the epigenetic modifier complexes that are recruited by this GFI1/LSD1 scaffold and act on chromatin structure and more specifically on histone modifications. Indeed, it was recently shown that GFI1B can recruit members of the so called BRAF-histone deacetylase (HDAC) (BHC) chromatin-remodelling complex that contains LSD1, CoREST, HDACs as well as a number of the High Mobility Group of proteins (HMG20A and -B) and other associated proteins (McClellan *et al*, 2019). The Nucleosome Remodeling and Deacetylase (NuRD) complex would be an excellent candidate as well, since similar to the BHC complex it also facilitate histone deacetylase mediated chromatin condensation and also contains LSD1 (Lai & Wade, 2011; Ramirez & Hagman, 2009; Wang *et al*, 2009).

The NuRD complex differs however from the BHC complex in that it contains seven different proteins divided into two sub-complexes, one which comprises an ATP-dependent nucleosome remodeling activity and another which harbors HDACs targeting H3K9 or H3K27 (Ahringer, 2000; Flanagan *et al*, 2005; Kelly & Cowley, 2013). Characteristic for the NuRD complex are its major components, the closely related proteins CHD4 (Chromodomain Helicase DNA Binding Protein 4 or Mi-2β), CHD3 and CHD5(Mills, 2017). CHD4 contains an SNF helicase domain and PhD/Chromo domains that mediate its interaction between nucleosomes and methylated histones(Woodage *et al*, 1997). The methyl-CpG-binding domain proteins MBD2 and MBD3 represent its non-enzymatic components and link the ATPase remodeling activities to the HDAC1 and -2 containing subcomplexes. The metastasis-associated proteins MTA1, MTA2, and MTA3 are also part of this subcomplex and mediate binding to DNA, to HDAC1, and to other transcription factors that can recruit NuRD to specific loci in the genome (Kumar *et al*, 2003; Ma *et al*, 2016; Yao & Yang, 2003). The proteins Rbbp7 and Rbbp4 bind histone H4 and most likely have roles as scaffolds(Kloet *et al*, 2015; Schmidberger *et al*, 2016). The GATA zinc-finger domain-containing proteins GATAD2A and –B directly interact with MBD2/3 and are also canonical members of the NuRD complex, although their precise function remains to be determined (Sharifi Tabar *et al*, 2019).

The NuRD complex can mediate both transcriptional repression or activation (Denslow & Wade, 2007; Lai & Wade, 2011) and can be recruited to sites of bivalent or “poised” targets in chromatin that are “primed” to be efficiently activated or repressed by modifying the histone marks during progenitor self-renewal or differentiation (Voigt *et al*, 2013; Zuo *et al*, 2009). Bivalent target promoters or genomic loci with enhancers show both repression marks such as H3K27me3 and activation marks such as H3K4me2 or H3K27ac at the same time and also feature modifications such as H3K4me1, which identify so called primed or induced enhancers (Voigt *et al*., 2013). To exert its function in a tissue and differentiation stage-specific manner, NuRD associates for instance during lymphoid development with lineage specific transcription factors and co-regulators(Denslow & Wade, 2007), such as IKZF1 (IKAROS), BCL6 or BLIMP1 (Dege & Hagman, 2014; Oestreich & Weinmann, 2011; Sridharan & Smale, 2007).

Here we show that GFI1 and GFI1B interact with members of the NuRD complex, notably with the chromodomain helicase DNA binding protein 4 (CHD4). In myeloid progenitors, GFI1 occupies chromatin together with CHD4 at specific target regions that are different from those regions occupied by GFI1 or CHD4 alone. These target regions bear characteristics of open chromatin and histone modifications that are associated with active transcription or poised enhancers. GFI1 occupies promoters and genomic sites that are different from those occupied by GFI1/CHD4 complexes and are upstream of different groups of genes that can be distinguished by the enrichment of either *Irf1* or *Spi1* DNA binding consensus sequences, respectively. During neutrophil differentiation, different levels of chromatin closure and a reduction of H3K4me levels is seen depending on whether sites are occupied by GFI1, CHD4 or GFI1/CHD4 complexes. Lastly, GFI1, CHD4 or GFI1/CHD4 occupy promoters of genes that fall into three distinct groups termed “immune system”, “chromatin/nucleosome assembly” and “metabolic process”, respectively, that can be both up- and downregulate during neutrophil differentiation.

## RESULTS

### GFI1 associates with the nucleosome-remodeling and histone-deacetylase (NuRD) complex

We had used AP-MS (affinity purification and mass spectrometry) to identify proteins that co-purified with Flag-tagged versions of GFI1 and GFI1B in HEK293 cells(Shooshtarizadeh *et al*, 2019; Vadnais *et al*, 2018). This approach revealed the presence of members of the NuRD complex such as Chromodomain Helicase DNA Binding Proteins 3 and -4 (CHD3 and -4), Metastasis Associated 1, and – 2 (MTA1 and -2) and the chromatin remodeling factor Retinoblastoma Binding Protein 4 (RBBP4, also called Chromatin Assembly Factor 1 or CAF-1) in both isolated GFI1 and GFI1B complexes (Fig. 1A, EV Fig. 1A). The peptide coverage for these factors was in a similar range as for those proteins that are known to associate with GFI1 and GFI1B such as HDAC1 and members of the CoREST complex such as RCOR1, 2 or -3 (Fig. 1A). To validate these findings, we used a BioID approach in HEK293 cells for GFI1 associated proteins, compiled the data with known interactions from the IntAct and BioGrid databases(Orchard *et al*, 2014; Oughtred *et al*, 2019) and observed that GFI1 has the potential to interact with four major complexes, notably with the NuRD complex, in agreement with our findings from the AP-MS experiment, but also with the SWI/SNF, CtBP and Cohesin complexes (Fig. 1B, suppl. Table 1).

**Figure 1:**
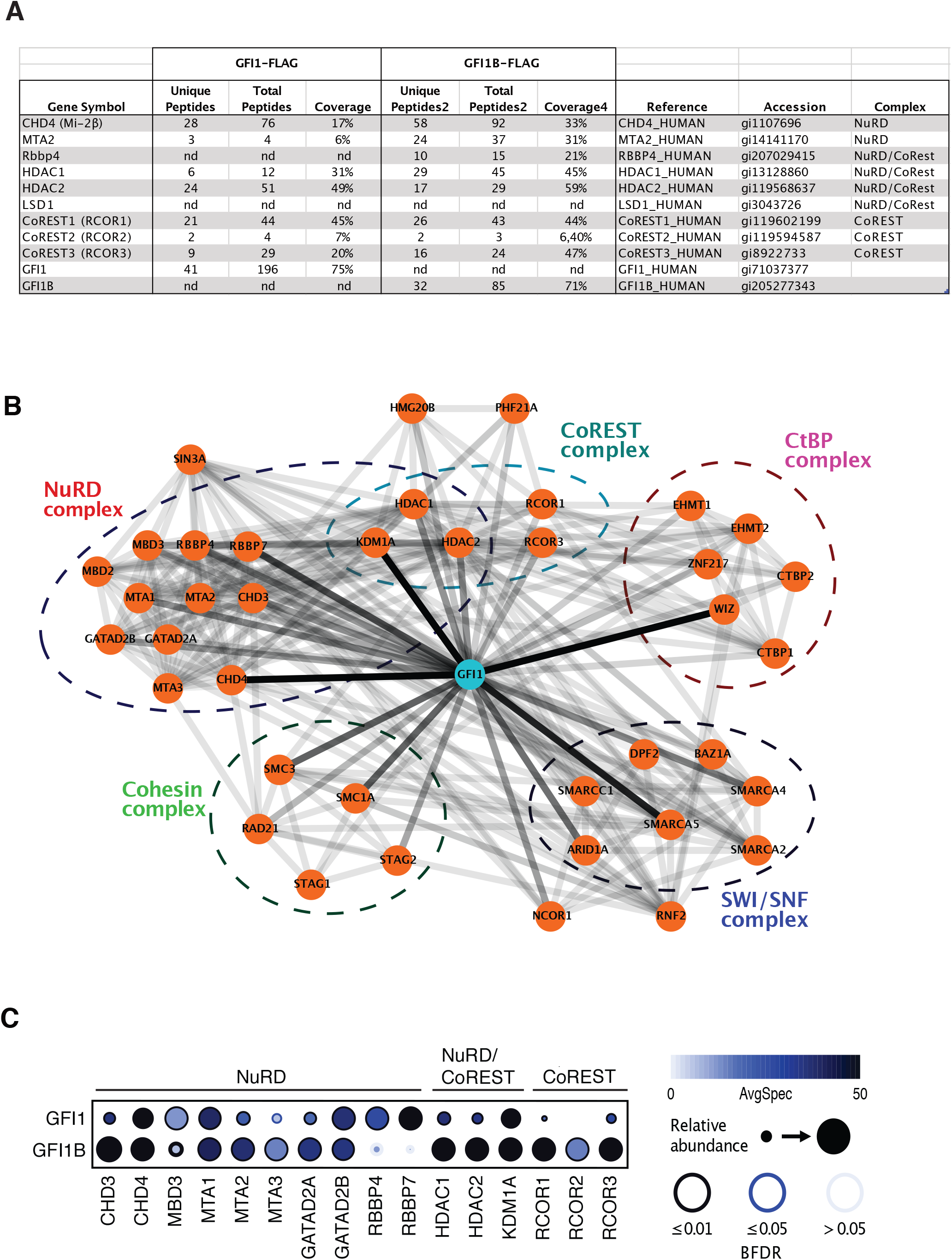
GFI1 interacts with protein components of the NuRD complex. **A)** Result from AP-MS experiments with GFI1- or GFI1B Flag expressed in 293T cells for NuRD and Co-Rest complexes. **B)** Interaction map of the GFI1 proteome obtained through BioID data. The interactions between GFI1 and the preys are ’’ weighted ’’ in color according to AvgSpec (e.g.: GFI1-KDM1A: 104 AvgSpec, darker color; GFI1-HDAC2: 44 AvgSpec less dark color). The known Prey-Prey interactions (e.g., KDM1A-HDAC1) were imported from databases IntAct and BioGrid, and are represent by less dark connections, since they cannot be normalized to the GFI1-BioID screen. GFI1-prey interactions remain(Voigt *et al*., 2013)visible. **C)** Dot Plot showing BioID interactions of GFI1-BirA*-Flag or GFI1B-BirA*-Flag with the indicated members of the NuRD or CoREST protein complexes. The node color depicts the average spectral counts. The relative abundance of prey versus the bait is shown by the circle size. The edge color represents the confidence score of the BioID/MS interaction (5% < BFDR as low confidence score, 1% < BFDR ≤ 5% as medium confidence or BFDR ≤ 1% as high confidence).

Next, we compared the data with our previously reported BioID experiment for GFI1B(Shooshtarizadeh *et al*., 2019) and found again members of the NuRD complex as high ranking candidates for new binding partners of both GFI1 and GFI1B (Fig. 1B, supp. Table 1), and proteins already known to bind to these two factors such as HDAC1 and -2, LSD1 and the CoREST proteins RCOR1, -2 and -3 (Fig. 1C, suppl. Table 1). Quantification of the mass spectrometry results showed high peptide coverage of CHD3 and CHD4, MTA1, -2, -3, and GATA Zinc Finger Domain Containing 2A and 2B (GATAD2A and GATAD2B); the abundance of recovered peptides for GFI1B being higher than for GFI1 (Fig. 1C). Also detectable were peptides for other NuRD complex members such as the Methyl-CpG Binding Domain Protein 3 (MBD3) and RBBP4 and - 7 (Fig. 1C, suppl. Table 1).

### NuRD components CHD4 and MTA2 associate with GFI1 C- and N-termini

Next, we expressed Flag-tagged versions of GFI1 and GFI1B in HEK293, precipitated nuclear extracts with anti-Flag agarose and analyzed the collected proteins per western blot. We were able to detect CHD4, MTA2, RBBP4/6 and, as positive controls, HDAC1 and LSD1 with both GFI1 and GFI1B (Fig. 2A). Also, samples from THP-1 cell extracts incubated with an anti GFI1 antibody contained both CHD4 and MTA2 protein (Fig. 2B, upper panel). Conversely, an anti MTA antibody precipitated both GFI1 and CHD4 in THP-1 cells (Fig. 2B lower panel) demonstrating that both CHD4 and MTA2 can interact with GFI1 at endogenous expression levels (Fig. 2B). In addition, Flag-tagged GFI1 and MTA2 showed similar interactions in the presence of absence of Ethidium-bromide or Benzonase (EV Fig. 1B), indicating that this interaction is independent of the presence of DNA.

**Figure 2:**
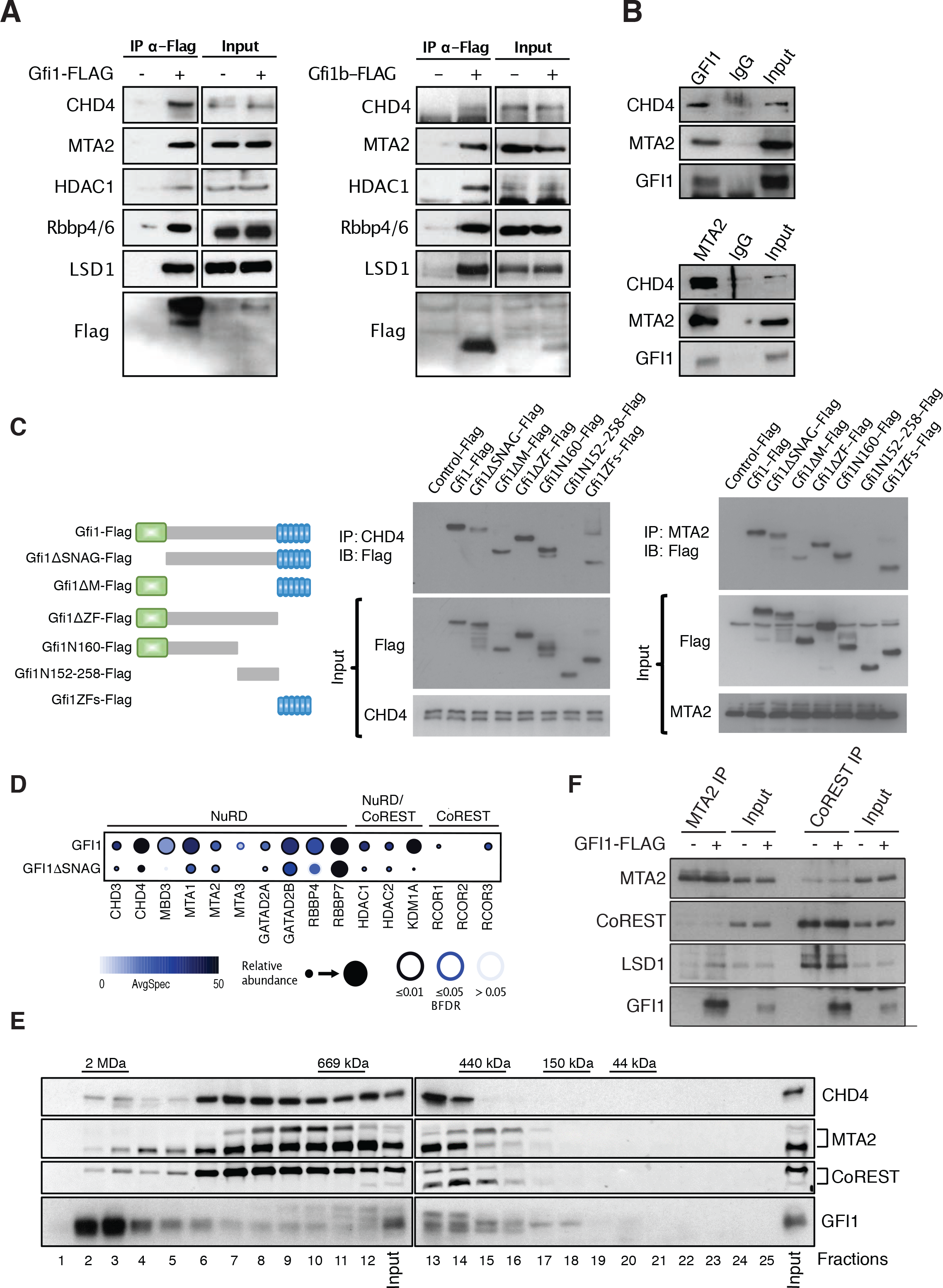
Association of GFI1 with NuRD complex requires the SNAG domain. **A)** GFI1-Flag fusion protein was immunoprecipitated from 293T cells; precipitates were separated by SDS–PAGE and blotted for the indicated proteins. **B)** Immunoprecipitation with anti GFI1 or anti MTA2 antibodies from THP-1 cell extracts. Precipitates were analyzed by Western blot for the presence of CHD4, MTA2 or GFI1. **C)** Schema of different GFI1-Flag fusion proteins that were expressed in HEK293T cells. Extracts of transfected cells were precipitated with anti CHD4 or MTA2 antibodies, separated by SDS-PAGE and analyzed by Western blotting with an anti-Flag antibody. **D)** Dot Plot showing BioID interactions of GFI1-BirA*-Flag or GFI1 lacking the SNAG domain (GFIΔSNAG-BirA*-Flag) with the indicated members of the NuRD or CoREST protein complexes. The node color depicts the average spectral counts. The relative abundance of prey versus the bait is shown by the circle size. The edge color represents the confidence score of the BioID/MS interaction (5% < BFDR as low confidence score, 1% < BFDR ≤ 5% as medium confidence or BFDR ≤ 1% as high confidence). **E)** Nuclear extracts from Kasumi 1 cells were fractionated using a Superose 6 10/300GL column; 0.5 ml fractions were collected, TCA-precipitated and loaded on SDS-PAGE for western blot analysis; immunoblots were probed with the indicated antibodies. **F)** HEK293T cells were transfected to express a full-length Flag-tagged GFI1 protein. Extracts were immunoprecipitated with anti-MTA2 or anti-Co-Rest antibodies and the precipitates were analyzed by Western blot for the presence of MTA2, Co-Rest, LSD1 or GFI1.

To determine the region of interaction between GFI1 and CHD4 or MTA2, we transfected different Flag-tagged truncation- and deletion mutants of GFI1 into HEK293 cells and precipitated extracts with anti-CHD4 or anti MTA2 antibodies. Analysis of the collected proteins by Western blot revealed that deletion of the SNAG domain weakened the interaction with either CHD4 or MTA2 (Fig. 2C) and that the presence of the SNAG- and the zinc finger domains alone could maintain an interaction with these two proteins (Fig. 2C). A new BioID experiment with either the full length GFI1 or a truncated version GFI1 lacking the 20 aa N-terminal SNAG domain (GFI1ΔSNAG) as baits confirmed, as previously reported(Saleque *et al*., 2007), that the interaction of GFI1 with the LSD1/CoREST complex requires the SNAG domain (Fig. 2D, EV Fig. 1C, suppl. Table 1). The experiment also showed that the interaction of GFI1 with members of the NuRD complex such as CHD3, -4, MBD3, MTA3, GATAD2A and HDAC1 and -2 seemed to be dependent on the presence of the SNAG domain. In contrast, RBBP7 or GATAD2B do not seem to require the SNAG domain for interaction with GFI1 (Fig. 2D), suggesting that their binding to GFI1 may be indirect via other members of the NuRD complex. A GO term analysis indicated that biological processes including nucleosome disassembly and protein acetylation and the molecular functions transcription coregulator and lysine acetylated histone binding were lost in the BioID with GFI1ΔSNAG (EV Fig. 1D, E, suppl. Table 2).

The majority of total nuclear GFI1 is eluted by size exclusion chromatography (SEC) in two complexes at around 2 MDa and 0.5 MDa from a Kasumi cell extract (Fig. 2E). This indicated that GFI1 associates with large complexes, such as the NuRD and Co-REST complexes as indicated by the presence of CHD4, MTA2 and CoREST at similar exclusion sizes of 0.5MDa (Fig. 2E). Both GFI1 and LSD1 but not CoREST were enriched in extracts precipitated anti MTA2 antibodies from HEK293 cells expressing a Flag-tagged version of GFI1 (Fig. 2F). In the absence of GFI1, anti MTA2 antibodies could still precipitate LSD1, albeit at lower levels (Fig. 2F). Anti-Co-REST antibodies precipitated both GFI1 and LSD1, but to a much a lesser extent MTA2 regardless of whether GFI1 was expressed or not (Fig. 2F). This suggests that GFI1 may interact with components of the NuRD complex such as MTA2 independently from the LSD1/CoREST complex.

### GFI1 and CHD4 co-occupy sites of open chromatin and active transcription

We chose primary murine GMPs, in which GFI1 is expressed(Zeng *et al*, 2004) to further investigate the association between NuRD and GFI1. ChIP-seq experiments with antibodies against murine GFI1 and CHD4 in wild type (WT) GMPs showed 3188 peaks for GFI1 and 4236 peaks for CHD4 and revealed that at 1128 sites occupation of GFI1 and CHD4 overlapped (Fig. 3A). The sites that are co-occupied by GFI1 and CHD4 are primarily at promoter- (∼30%) and intergenic regions (∼38%) (Fig. 3B). When we compared the overall binding of CHD4 to chromatin in sorted GMPs from WT with gene deficient (*Gfi1*^-/-^) mice, we observed that of the 1128 sites that were co-occupied by both GFI1 and CHD4 in WT cells, 841 lost enrichment of CHD4 when GFI1 was absent (Fig. 3B-D), most of them in regions <3kb from the TSS. At the remaining 287 sites, CHD4 was still enriched regardless of GFI1’s presence or absence (Fig. 3C, D). Except for 9 new CHD4 enriched sites, no significant new gain in CHD4 bound sites was observed in *Gfi1^-/-^* GMPs (Fig. 3C). The 3108 sites that were occupied only by CHD4 in WT cells, remained largely unaffected by *Gfi1* deficiency, i.e., CHD4 remained at these sites (Fig. 3C).

**Figure 3:**
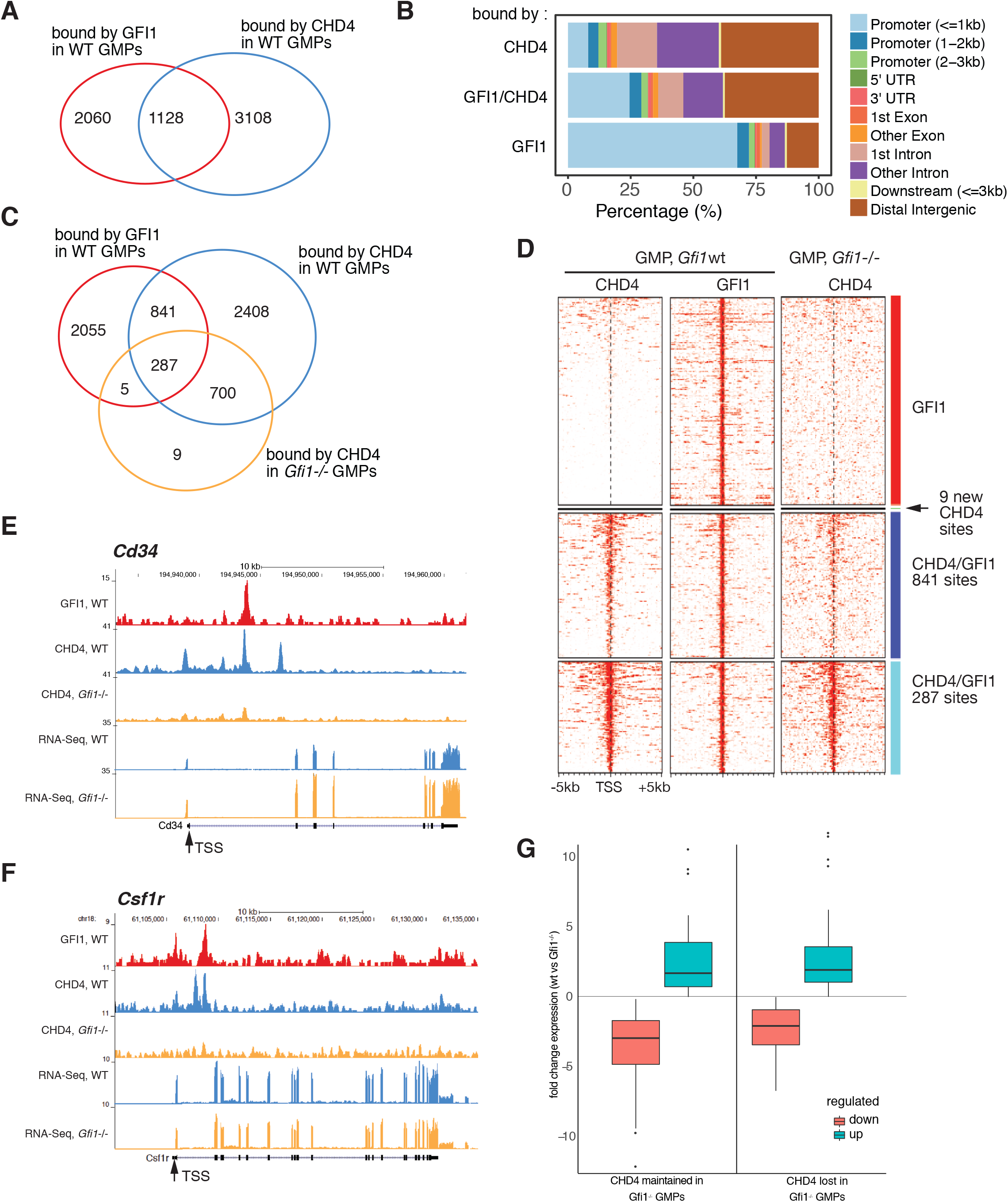
GFI1 and NuRD complex member CHD4 co-occupy sites at chromatin from granulocyte myeloid progenitors (GMPs) **A)** Venn diagram indicating the number of sites occupied by GFI1, CHD4 or both in GMPs according to ChIP-seq experiments done with GMPs. **B)** Relative distribution of sites occupied by GFI1, CHD4 or both in percent of all sites in promoter regions, 3’ and 5’ untranslated regions (UTRs), introns, exons or other distal regions. **C)** Venn diagram indicating the number of sites occupied by GFI1 in WT GMPs, CHD4 in WT GMPs and CHD4 in GFI1 deficient GMPs. **D)** Heat map of ChIP-seq results for genes occupied by GFI1, or GFI1 and CHD4 in WT GMPs or in GMPs sorted from GFI1 deficient mice (GFI1 KO). Genes are sorted according to GFI1 occupation. Red: genes occupied by GFI1, but not by CHD4, dark blue: genes occupied by GFI1 and CHD4, which lose CHD4 occupation in GFI1 KO cells, pale blue: genes occupied by GFI1 and CHD4, which maintain CHD4 occupation in GFI1 KO cells **E)** Schematic depiction of the locus encoding murine CD34 and **F)** the *Csf1r* locus encoding M-CSFR. Shown is the enrichment of reads after ChIP-seq with antibodies against GFI1 or CHD4 in GMPs from either WT or GFI1KO mice and the enrichment of reads after an RNA-seq experiment from WT of GFI1 KO GMPs. The transcription start site is indicated (TSS). **G)** The promoters of the genes targeted by GFI1 and by CHD4 were separated into two groups according to the presence of CHD4 in *Gfi1* KO cells (w_chd4 and wo_chd4. The groups were then separated into two subgroups according to the direction of the fold change in expression after *Gfi1* deletion (up or down). For the comparison w_chd4_up vs wo_chd4_up, the p-value is 0.80. For the comparison w_chd4_down vs wo_chd4_down, the p-value is 0.0095.

Examples for loci occupied by GFI1/CHD4 are the genes *Cd34*, *Csf1r* (encoding M-CSFR) or *Csf1* (encoding M-CSF) (Fig. 3E, EV Fig. 2A). *Gfi1* deletion is associated with a loss of CHD4 at these sites and RNA-seq data showed that this leads to the upregulation of *Cd34* mRNA (Fig. 3E) but does not affect *Csf1r* and *Csf1* mRNA levels (Fig. 3F and EV Fig. 2A). For further validation, we chose five other genes selected from the group of 841 loci where CHD4 binding was lost in the absence of GFI1 according to the ChIP-seq data. ChIP-qPCR on these gene loci for CHD4 in primary WT and *Gfi1^-/-^* GMPs showed loss of CHD4 enrichment confirming the results of the CHD4 ChIP-seq experiment (EV Fig.2B). RNA-seq and flow cytometric analysis demonstrated that *Chd4* mRNA and CHD4 protein levels were not altered in *Gfi1^-/-^* GMPs compared to WT GMPs (EV Fig. 2C, D). Genes that lost CHD4 binding at their promoters in the absence of GFI1 or maintained CHD4 did not show a significantly different up- or downregulation of gene expression in *Gfi1^-/-^*versus wt GMPs (Fig. 3G). Also, up-or downregulation of GFI1 target gens in *Gfi1^-/-^*versus wt GMPs did not correlate with presence of absence of CHD4 (EV Fig. 3A, B). This suggested that the deletion of GFI1 affects gene expression independently of the presence of CHD4.

To test whether GFI1/CHD4 complexes affect chromatin remodeling, we performed ATAC-seq and ChIP-seq analyses of GMPs with antibodies against methylated and acetylated histone H3. The obtained data were filtered for promoter regions defined here as regions located between less than 2kb upstream and less than 500 bp downstream of transcription start sites (TSS), and separately also for enhancer regions selected based on the criteria defined in the Fantom5 enhancer atlas(Andersson *et al*, 2014). The data were ordered according to CHD4 occupation and were separated into three groups: occupation by CHD4 alone, by GFI1/CHD4 together or by GFI1 alone (Fig. 4A). We observed that promoters occupied by GFI1 show higher levels of markers associated with active transcription such as H3K4me3 and H3K27ac than the promoters occupied by CHD4 alone. Promoters co-occupied by both GFI1 and CHD4 showed intermediate levels of H3K4me3 and H3K27ac (Fig. 4B). In contrast, promoters occupied by CHD4 showed higher levels of H3K27me3 H3K9me3 and H3K4me2, all associated with transcriptional repression, than promoters occupied by GFI1 or by both GFI1 and CHD4 (Fig. 4B), with a similar pattern for the levels of H3K4me2 and H3K4me3, which are markers of transcriptional activation (Fig. 4B). This situation is exemplified by the promoter regions of the genes encoding the GFI1 targets *Csf1* (Fig. 4C) and *Csf1r* (EV Fig. 4), which are both expressed in GMPs (Fig. 3E, F).

**Figure 4:**
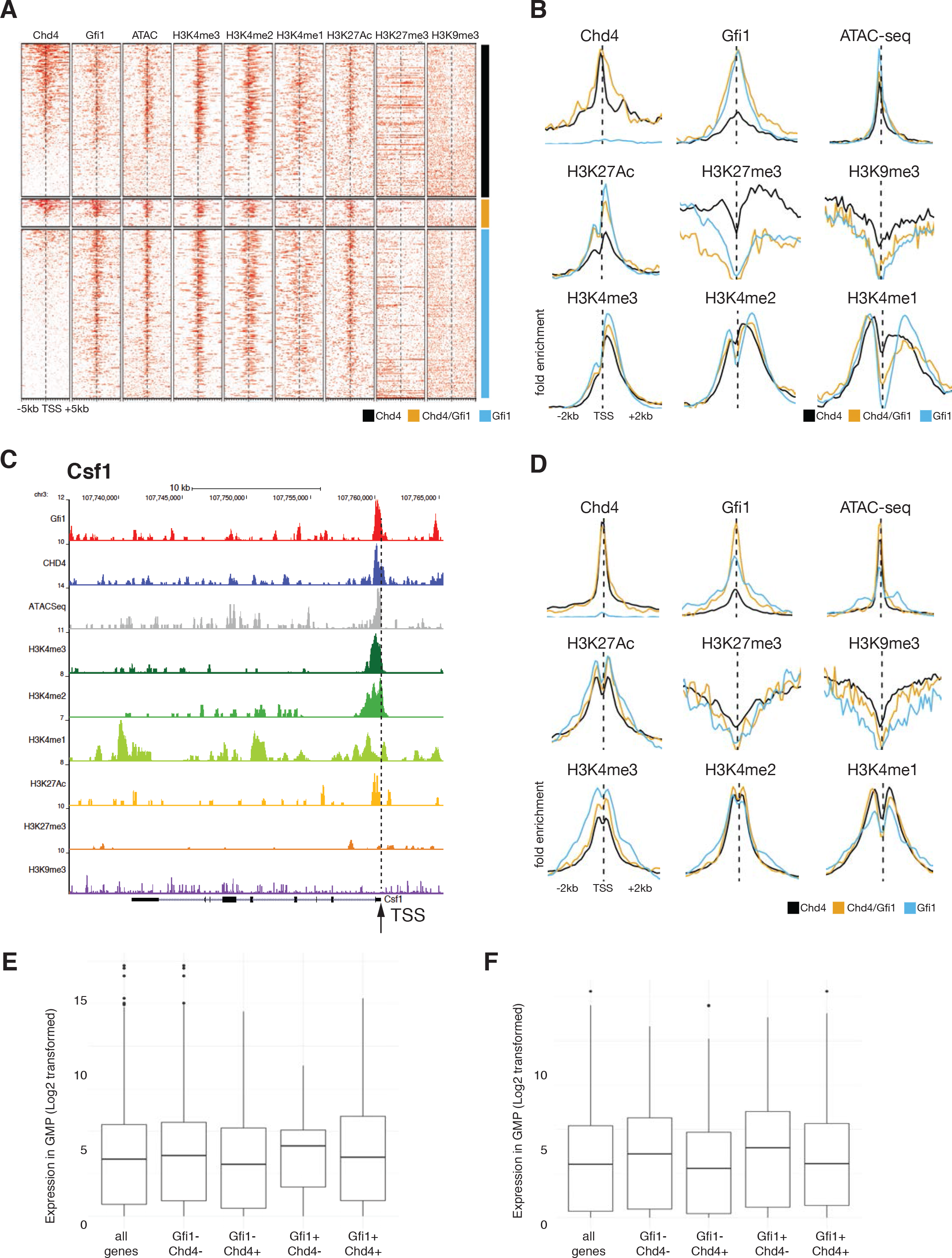
GFI1 and GFI1/CHD4 occupy actively transcribed regions. **A)** Heatmap of ATAC-seq and ChIP-seq analyses obtained with antibodies against GFI1, CHD4 and methylated and acetylated histone H3 from GMPs. Shown are reads for promoter regions with 5kb 5’ and 3’ of TSS ordered according to CHD4 occupation and were separated into three groups: occupation by CHD4 alone, by GFI1 and CHD4 and by GFI1 alone. **B)** Aggregation plots for promoters occupied by GFI1, CHD4 and methylated and acetylated histone H3 with the data shown in A). Shown is the fold enrichment over a region of 2kb 3’ and 5’ of the TSS. **C)** Exemplary depiction of the ATAC-Seq and ChIP-Seq data at the GFI1 target gene *Csf1*. Indicated are the tracks corresponding to the individual experiments, the gene and the TSS. **D)** Aggregation plots for enhancers occupied by GFI1, CHD4 and methylated and acetylated histone H3. Shown is the fold enrichment over a region of 2kb 3’ and 5’ of the TSS. **E)** Analysis of RNA-seq data from GMPs of genes that are next to the promoters or **F)** enhancers and were occupied by GFI1, CHD4 or both CHD4 and GFI1.

H3K9 and H3K27 methylation and acetylation patterns at enhancers correlated similarly with GFI1 and CHD4 occupation to those at promoters (Fig. 4D). Also, we found relatively higher levels of H3K4me1 and lower levels of H3K4me3 at enhancers occupied by CHD4 only and a relative loss of H3K4me1 and gain of H3K4me3 at sites occupied by GFI1 only and values for GFI1/CHD4 occupation in between (Fig. 4D). Analysis of RNA-seq data from GMPs of genes that are next to the promoters or enhancers defined here and were occupied by GFI1 showed higher expression levels than the genes next to sites bound by CHD4 or both CHD4 and GFI1 (Fig. 4E and 4F, respectively). These findings indicate that GFI1/CHD4 complexes may occupy active or bivalent promoters and active enhancers that have a more open chromatin configuration when CHD4 is absent or have a more closed chromatin conformation when GFI1 is absent and suggest a repressive action is mediated by CHD4 rather than by GFI1.

### De novo motif analysis highlights myeloid-specific gene network

Next, we compared our ChIP-seq data for GFI1 and CHD4 binding and H3K4 methylation with a data set for the transcription factor CEBPα, also done in GMP cells, which has been shown to co-occupy promoter sites together with GFI1 in myeloid cells(Pundhir *et al*, 2018). The data were again analyzed for promoter regions as defined above, they were ordered according to GFI1 binding and were separated into four groups: occupation by GFI1 alone, by GFI1/CEBPα, by GFI1/CEBPα/CHD4 and by GFI1/CHD4 (Fig. 5A). A *de novo* motif analysis(McLeay & Bailey, 2010) of consensus DNA binding sites at promoter regions revealed that the loci bound by GFI1 or by GFI1/CEBPα were very highly enriched for the GFI1/GFI1B DNA binding motif, as expected, but the enrichment was even higher for the consensus DNA binding motif for the transcription factor IRF1, which contains the 5’-AANNGAAA-3’ core sequence for all IRF factors (Fig. 5B). In addition, *Ets2*, *Stat3* and *Sox3* binding motifs were only present at sites occupied by GFI1 and *Klf*, *Spi1*, *Runx* and *Atf3* binding motifs were only present at sites occupied by GFI1/CEBPα (Fig. 5B). At sites where GFI1 and CHD4 are both present, neither *Irf* nor *Gfi1* consensus sites were enriched. However, the binding motifs for SPI1 (PU.1) and the related factor SPI-C were found to be enriched with highest E values (Fig. 5C). Sites occupied by GFI1/CHD4/CEBPα showed again enrichment for the PU.1 and SPI-C motifs, but in addition also the binding sequences for CEBPα and RUNX1 (Fig. 5B). This differential enrichment of binding motifs suggests that GFI1 or GFI1/CEBPα complexes bind to other genomic loci that those occupied by GFI1/CHD4 or GFI1/CEBPα/CHD4 complexes. Aggregation plots showed that H3K4me2 and H3K4me1 methylation levels at promoter regions occupied by GFI1 or GFI1/CEBPα complexes are differentially depleted or enriched at the TSS or 3’ of the TSS, respectively, according the to presence of CHD4 (Fig. 5C). The data suggest that GFI1 is more efficient in the depletion of H3K4me2 and -me1 marks in the presence of CHD4 (Fig. 5C). This also suggests that GFI1 and CEBPα are associated with promoters with a higher transcriptional activity than those where CHD4 is also present either together with GFI1 or GFI1/ CEBPα complexes, and, in addition, suggesting again that the presence of CHD4 is associated with repressive histone marks.

**Fig. 5.**
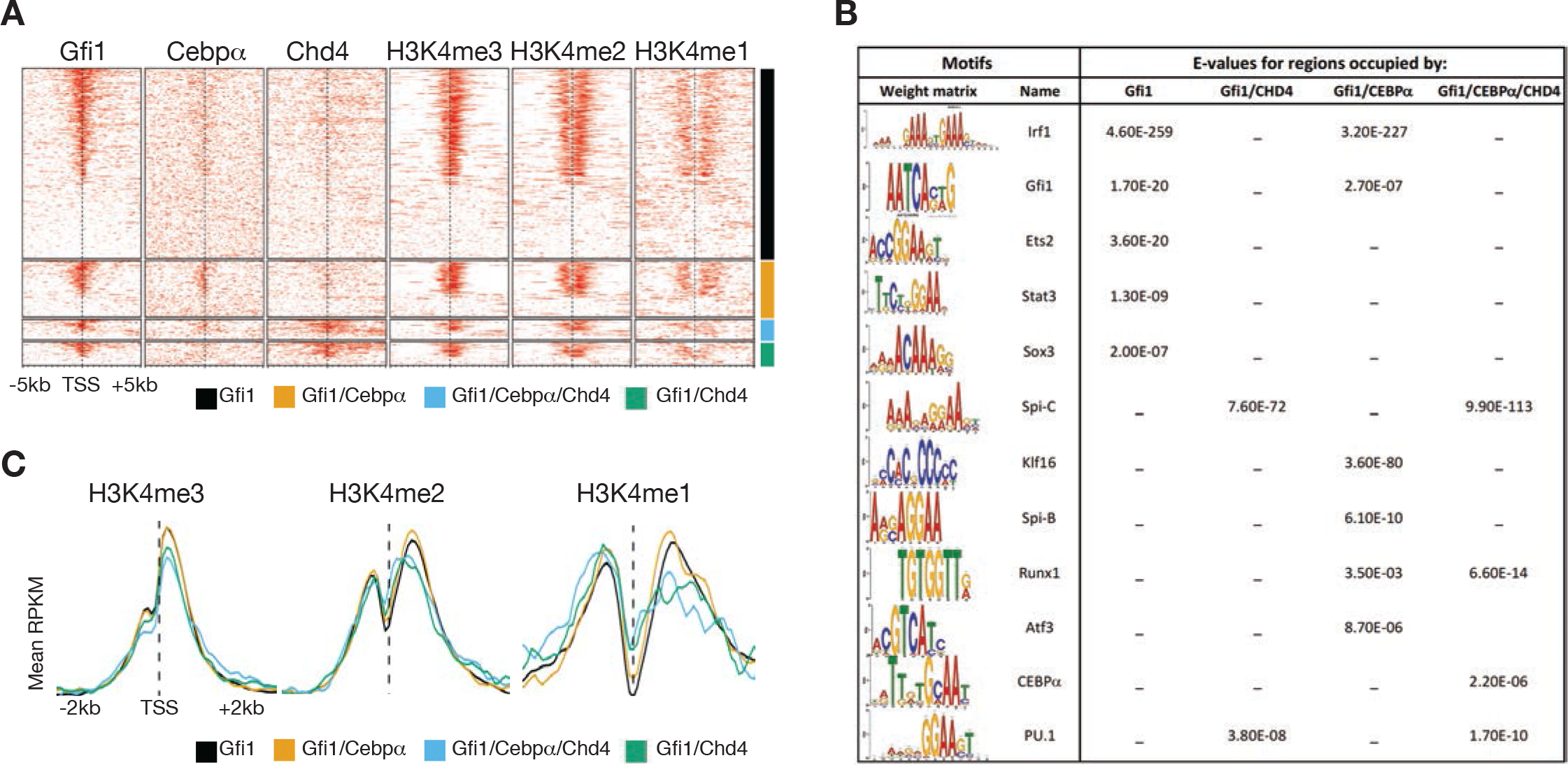
De novo motif analysis and histone methylation aggregation plots for sites occupied by GFI1, CHD4 and CEBPα in GMPs. **A)** Comparison of ChIP-seq data for the occupation of GFI1, CHD4 and H3K4 methylation at promoter regions in GMPs with an analogous ChIP-seq data set for the transcription factor CEBPα. Data are ordered according to GFI1 occupation and are separated into four groups: occupation by GFI1 alone, by GFI1/CEBPα, by GFI1/CEBPα/CHD4 and by GFI1/CHD4. **B)** D*e novo* motif analysis of consensus DNA binding sites at promoter regions by GFI1 alone, by GFI1/CEBPα, by GFI1/CEBPα/CHD4 and by GFI1/CHD4. **C)** Aggregation plots of histone H3K4 methylation at promoters as defined in **A)** for regions 2kb 5’ or 3, of the TSS.

### Chromatin remodeling by GFI1, CHD4 or both CHD4 and GFI1 during neutrophil differentiation

The developmental steps from GMPs to mature neutrophils have been clarified using surface markers to define pre-neutrophils (preNeu), immature neutrophils (immNeu) and mature neutrophils (matNeu) stages (Evrard *et al*, 2018). To explore how sites that are occupied by GFI1, CHD4 or both in GMPs are altered during neutrophil differentiation with regard to chromatin openness or H3K4 dimethylation levels, we sorted GMP, preNeu and matNeu populations from bone marrow following a published strategy(Evrard *et al*., 2018) (EV Fig. 5A). Monitoring GFP expression in these same subsets isolated from *Gfi1*^GFP^ knockin mice (Yucel *et al*, 2004) showed that the *Gfi1* gene is expressed and the promoter is active (suppl Fig. 5B). However, analysis by western blot of the sorted cells showed that GFI1 protein expression levels are higher in matNeus than in preNeu or immatNeu cells (EV Fig. 5C). RNA-seq reads of groups of genes specific for chemotaxis or phagocytosis were obtained and were congruent with the published pattern (EV Fig. 6A, B), similar to other genes regulated during myeloid differentiation indicating that the sorted populations represent indeed preNeu and matNeu cells as previously reported(Evrard *et al*., 2018) (EV Fig.7).

Next, we performed both ATAC-Seq and ChIP-Seq experiments to determine whether and how chromatin openness and H3K4 di-methylation levels change at the loci that are occupied by GFI1 and CHD4 or both in GMPs during neutrophil differentiation. Data were filtered as described above for promoter and enhancer regions, were ordered according to CHD4 occupation, and were separated into three groups: occupation in GMPs by CHD4 alone, by GFI1/CHD4 and by GFI1 alone (Fig. 6A, B). We compiled the data for the locus of a typical myeloid specific enhancer localized in the 3’ region of the *PLBD1* gene(Pundhir *et al*., 2018) and several sites downstream (Fig. 6C). Sites 3’ of the *Plbd1* gene that are occupied by GFI1/CHD4 in GMPs showed a decrease in ATAC-seq reads and a low level H3K4me2 marks compared to the site within the *Plbd1* gene where CHD4 is present without GFI1. At this site, ATAC-seq levels increased and H3K4me2 levels accumulated in preNeu and matNeu cells (Fig. 6C).

**Fig. 6:**
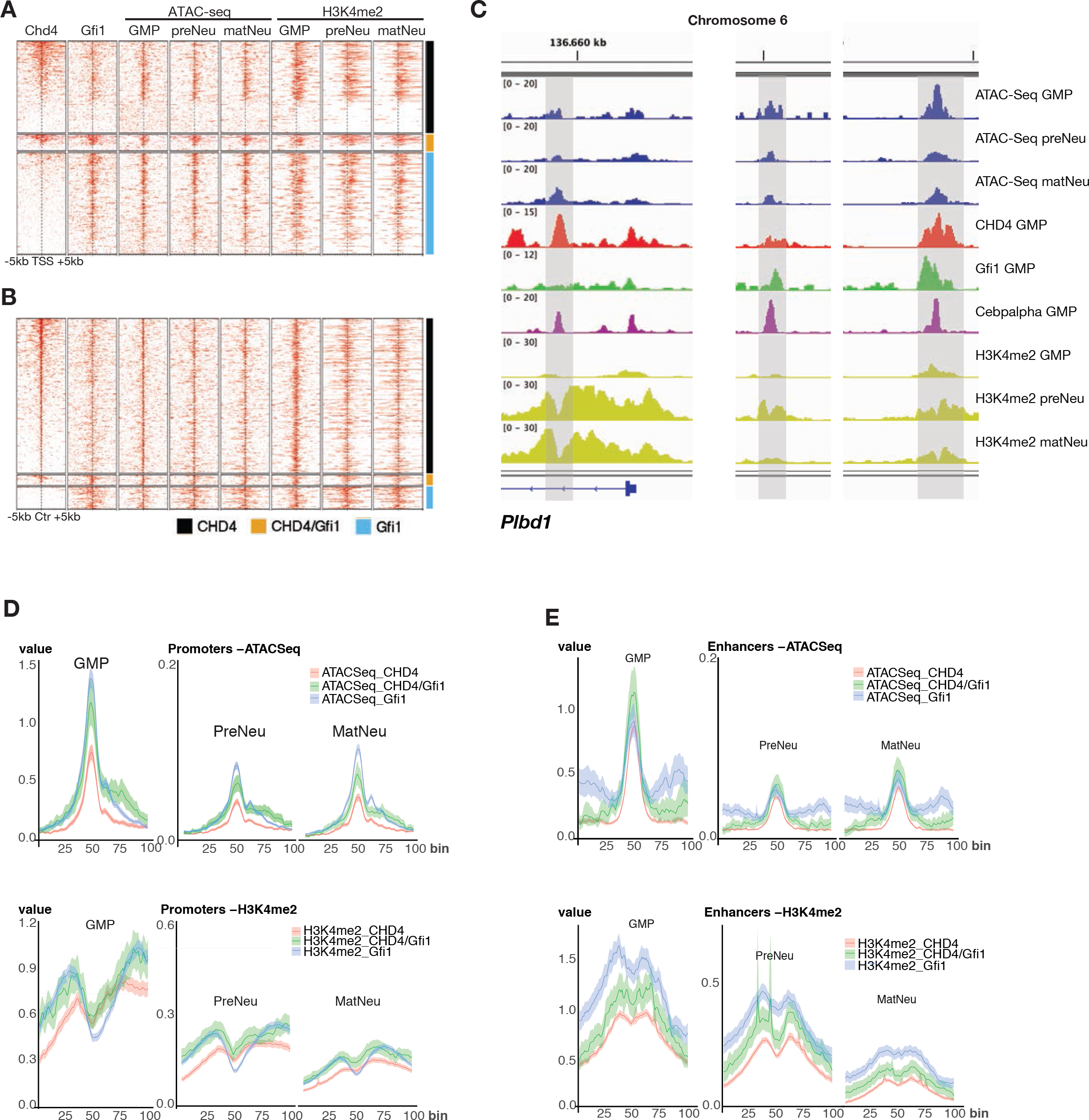
H3K4 methylation and chromatin status at sites occupied by GFI1, CHD4 or both CHD4 and GFI1 during neutrophil differentiation. **A)** Heat map of data from ChIP-Seq and ATAC-Seq experiments and to determine occupation of GFI1, CHD4 and H3K4 di-methylation patterns during neutrophil differentiation from GMPs via preNeu (preneutrophils) to matNeu (mature neutrophils). Data were filtered for promoter regions, ordered according to CHD4 occupation, and were separated into three groups: occupation by CHD4 alone, by GFI1/CHD4 and by GFI1 alone. **B)** Heat map of data from ChIP-Seq and ATAC-Seq experiments as in **A)**, but for enhancer regions. **C)** ATAC-Seq and ChIP-seq reads on loci covering the myeloid specific enhancers at the 3’ end of the *Plbd1* gene and at downstream regions for sites occupied by GFI1, CHD4, or both, CEBPα and carrying the indicated histone marks. **D)** Metagene analysis for ATAC-seq reads and H3K4me2 levels (**E**) in RPM values for GMPs, preNeu and matNeu cells at promoters occupied by CHD4 alone, by GFI1/CHD4 and by GFI1 alone. Shown are regions 2kb 5’ and 3’ of the TSS. **F)** Metagene analysis for ATAC-seq reads and for H3K4me2 levels (**G**) in mean RKPB for GMPs, preNeu and matNeu cells at enhancers occupied by CHD4 alone, by GFI1/CHD4 and by GFI1 alone. Shown are regions 2kb 5’ and 3’ of the site of the enhancer (Ctr).

To better quantify and facilitate the integration of signals from GMPs, preNeu und matNeu cells of regions that are occupied by GFI1, CHD4 or both GFI1 and CHD4 in GMPs, we performed a Metagene analysis(Joly Beauparlant *et al*, 2016). This allowed us to directly compare the enrichment profiles of H3K4me2 and ATAC-seq signals between cell types and experiments and to visualize the result. We used the same definition of promoter and enhancer regions as in the previous analyses to extract the enrichment signal, which was then normalized using the NCIS algorithm(Liang & Keles, 2012). This permitted us to compare the mean coverage values between cell types (GMP, preNeu, matNeu), since the normalization integrates both background noise and the size of the library (RPM). Bins cover 1000 base pairs 5’ and 3’ of the TSS for the promoter region and of the defined center region (Ctr) of the enhancer (at 50 bins).

Using the ChIP-seq data from GMPs, we compiled values from 2190 promoter sites occupied by GFI1, 1779 sites occupied by CHD4 and 341 sites occupied by both CHD4 and GFI1. We observed that mean coverage of ATAC-seq RPM values for these promoters were around 15- fold lower in preNeu and matNeu cells than in GMPs (Fig. 6D). Also, within preNeu and matNeu cells, they were significantly lower at sites that were occupied by CHD4 than those occupied by GFI1 in GMPs (p < 10^-30^, red lines Fig. 6D, suppl. Table 3), indicating that chromatin regions close during neutrophil differentiation and that sites occupied in GMPs by CHD4 are more contracted than sites occupied in GMPs by GFI1 or GFI1/CHD4. Similarly, at the same sites the mean coverage of H3K4me2 levels were lower overall in preNeu and matNeu cells compared to GMPs (Fig. 6D). They were also lower at the promoter sites occupied by GFI1 compared to sites occupied by CHD4 or GFI1/CHD4 complexes at the TSS (blue line Fig. 6D, supp. Table 2), suggesting that GFI1 may be active in removing methyl groups from H3K4 in GMPs and preNeu cells and that this is modified when CHD4 is present together with GFI1.

For enhancers, we included values from 382 sites occupied by GFI1, 2375 sites occupied by CHD4 and 173 sites occupied by both CHD4 and GFI1 in GMPs. The RPM values of ATAC-seq reads indicated a highly significant chromatin contraction for these sites in preNeu and matNeu cells versus GMPs (Fig. 6E, suppl. Table 1), but no differences were seen in each cell type between enhancer centers that were occupied by GFI1, CHD4 or GFI1/CHD4 in GMPs, only at regions 5’ and 3’ of the center when occupied by GFI1 (Fig. 6E, suppl. Table 2). H3K4me2 values at these enhancers were again lower in preNeu and matNeu cells than in GMPs (Fig 6E, suppl. Table 2). However, sites at the enhancer centers occupied in GMPs by CHD4 had significantly lower H3K4me2 levels than sites occupied by GFI1 (Fig. 6E, red lines and blue lines, respectively), whereas sites occupied by GFI1/CHD4 had intermediate levels (Fig. 6E, green line, suppl. Table 2). This indicates that in GMPs GFI1 and CHD4 act differently on H3K4 methylation at enhancer sites than at promoters and that this differential pattern is retained in preNeu and matNeus cells.

### Genes occupied by CHD4, GFI1 or both belong to the different categories

Next, we reordered the Chip-Seq and ATAC-seq data once again according to CHD4 occupation at promoter and enhancer sites, but now separated them into 6 groups: occupation by CHD4 alone, by GFI1/CHD4 or by GFI1 alone and according whether these groups of genes were up or down regulated during the differentiation from GMPs to preNeu and matNeus (Fig. 7A, B). A GO classification showed that genes occupied by CHD4 alone are found in pathways typical for the immune and inflammatory response regardless of whether they were up- or downregulated during differentiation from GMPs to matNeu cells (Fig. 7B). Up and downregulated genes co-occupied by both CHD4 and GFI1 encode regulators of chromatin assembly and nucleosome organization, while genes occupied by GFI1 alone are involved in metabolic processes (Fig. EV. 7), suggesting that GFI1, CHD4 and the GFI1/CHD4 complex regulate separate defined groups of genes during neutrophil differentiation.

**Fig. 7:**
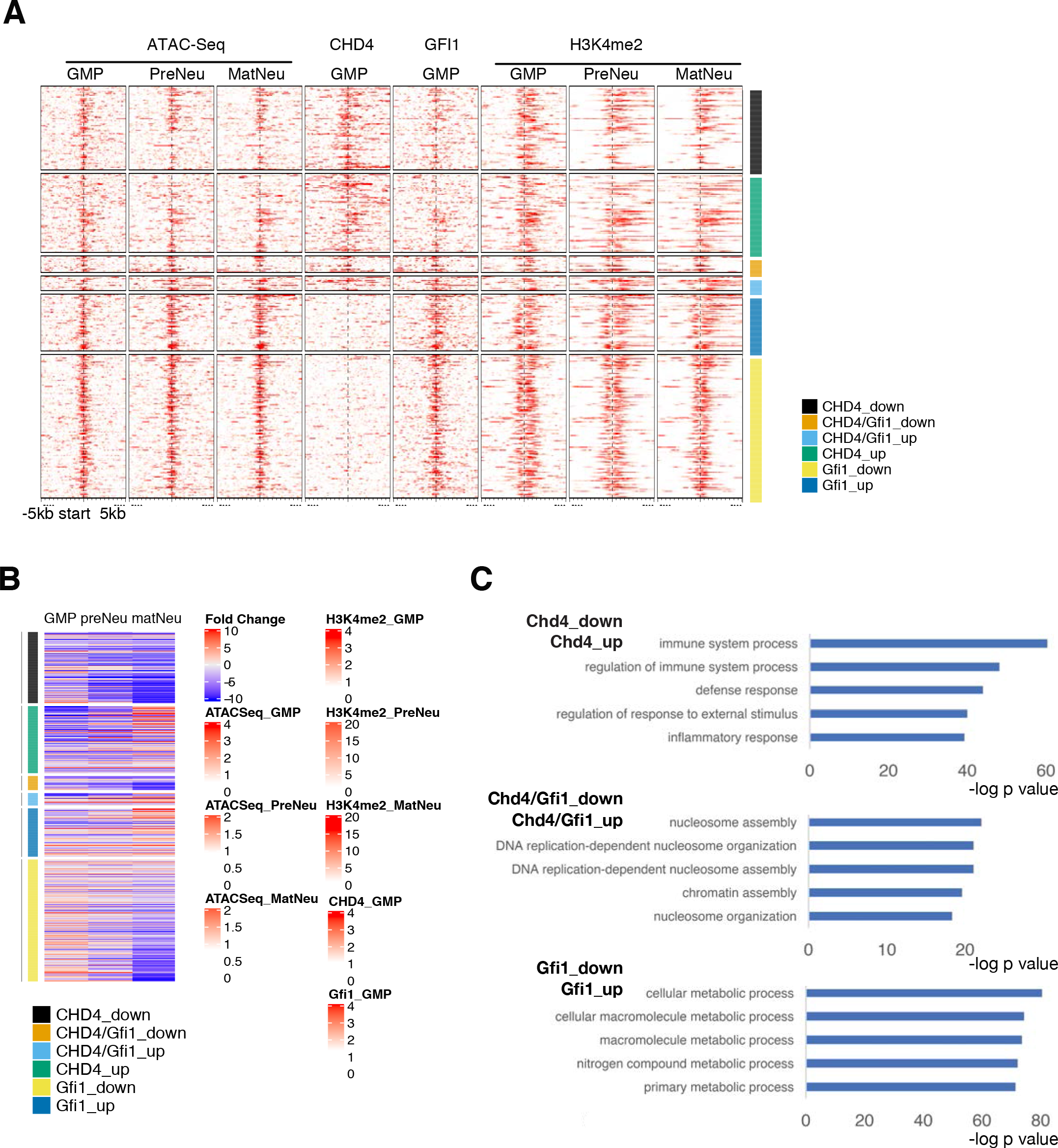
Chromatin status and H3K4 dimethylation pattern during neutrophil differentiation. **A)** Heat map of Chip-Seq and ATAC-seq data obtained in GMPs, preNeu and matNeu cells ordered according to CHD4 occupation at promoter sites in GMPs. The data were separated into 6 groups: occupation by CHD4 alone, by GFI1/CHD4 complexes or by GFI1 alone and according whether these groups of genes were up or down regulated during the differentiation from GMPs to preNeu and matNeu cells. **B)** Heat map of RNA-Seq data from GMPs, preNeu and matNeu cells, ordered according to genes with promoters occupied by CHD4 alone, by GFI1/CHD4 complexes or by GFI1 alone and according whether these groups of genes were up or down regulated during the differentiation from GMPs to preNeu and matNeu cells. **C)** GO classification of genes occupied by CHD4 alone, by GFI1/CHD4 complexes or by GFI1 alone and according whether these groups of genes were up or down regulated during the differentiation from GMPs to preNeu and matNeu cells.

## DISCUSSION

In the present study, we describe results of BioID experiments indicating that the SNAG domain and zinc-finger transcriptional repressor GFI1 associates with several chromatin remodeling complexes such as the NuRD complex, which also contains LSD1/CoREST, but also the CtBP and SWI/SNF chromatin remodeling and repressor complexes. We chose to elucidate the biological significance of the interaction with the NuRD complex, since we find that GFI1 can associate with almost all its components. We have focused on the chromodomain helicase CHD4, one of the NuRD complex components, and have used primary murine cells representing stages of neutrophil differentiation as a model system given the particularly important role of GFI1 in this process. We demonstrate that GFI1 can recruit CHD4 to specific sets of target genes that regulate processes such as nucleosome organization and chromatin assembly. While GFI1 occupies target gene promoters containing its own cognate DNA consensus sequence and those for IRF1, GFI1/CHD4 complexes target genes at consensus sites for ETS related factors such as SPI1 (PU.1) and SPIC. Analysis of histone modifications and chromatin structure indicates that GFI1 and GFI1/CHD4 complexes occupy active or bivalent promoters and active enhancers, both up and downregulated during neutrophil differentiation.

### Several regions of GFI1 take part in its association with components of NuRD complex

Previous studies showed that GFI1 binds to LSD1/CoREST and HDACs (Duan & Horwitz, 2003b; Saleque *et al*., 2007; Vassen *et al*, 2006), but these histone modifying enzymes are also members of the NuRD complex. It is thus possible that GFI1 interacts with the NuRD complex either via LSD1 or HDACs, or through CHD4 or MTA2, the proteins identified here to bind to GFI1. Immunoprecipitation and mass spectrometry data, BioID experiments with full length GFI1 and a GFI1 mutant that lacks the SNAG domain, biochemical analyses with mutated and truncated GFI1 proteins support a model in which GFI1 associates with the CHD4 and MTA2 components of the NuRD complex through several regions including the SNAG and zinc finger domains. Given that the SNAG domain specifically and directly interacts with LSD1(Lin *et al*., 2010; Saleque *et al*., 2007), it is possible that GFI1 acts as a bridging factor between LSD1/CoREST and components of the NuRD complex such as CHD4 and MTA2. SEC fractionation data suggest that these complexes are around 400-500 kDa, but also indicate that GFI1 is a member of other, high molecular weight complexes, in which NuRD components are less likely to be found. This would be in agreement with the notion that GFI1 acts as a member of several different chromatin remodeling complexes, which is also supported by our BioID results.

### GFI1 recruits the NuRD complex component CHD4 to active or bivalent promoters and active enhancers

Since we can demonstrate that GFI1 and CHD4 co-occupy promoter and enhancer sites in GMPs, it is conceivable that GFI1 recruits the NuRD complex to these regions in the chromatin. For a considerable fraction of these sites the recruitment is likely to occur directly through GFI1, since the deletion of GFI1 abrogates CHD4 occupation. The deletion of GFI1 did not lead to a large redistribution of CHD4 to new sites as it was observed in similar experiments with the transcription factor IKAROS in B lymphocytes(Oestreich & Weinmann, 2011), which underlines an important difference to our study with myeloid cells. According to the existing model, sites occupied by GFI1 should be depleted of H3K4me2, -me1 and H3K9acetyl marks and be transcriptionally silent (Olsson *et al*, 2016; Saleque *et al*., 2007). However, our ChIP-seq data suggest that GFI1 and CHD4 and GFI1/CHD4 complexes are located at transcriptionally active promoters in GMPs. The enrichment of markers for active transcription such as acetylation at H3K27 and di- and tri-methylation at H3K4 and the depletion of markers indicating transcriptional repression such as tri-methylation of H3K27 and H3K9 at sites where GFI1 binds support this notion. However, while H3K4me1 that indicates active transcription is strongly depleted at the TSS of GFI1 occupied genes, it is possible that these are rather bivalent promoters, which are characterized by the presence of both active and repressive histone marks.

Similarly, enhancers occupied by GFI1, CHD4 and GFI1/CHD4 complexes show H3K4me1 and H3K27 acetylation and absence or depletion of H3K27me3 or H3K9me3 marks, which is consistent with active enhancers as opposed to inactive or poised enhancers, which would be characterized by the presence of both H3K4me1 and H3K27me3. Of interest here is that sites occupied by CHD4 show relatively lower levels of active histone marks and relatively higher level of histone marks associated with inactive promoters and enhancers compared to sites occupied by GFI1 alone. Sites occupied by both GFI1 and CHD4 consistently show levels in between suggesting that GFI1 can modulate the effect of the NuRD complex on chromatin remodelling. RNA-seq data show that genes associated with promoters or enhancers occupied only by GFI1 are expressed at a higher level than genes occupied only by CHD4, while genes associated with promoters or enhancers occupied by both GFI1 and CHD4 have expression levels that fall in between these extremes. This supports the view that GFI1 is associated with active promoters and enhancers and can modulate the effect of the NuRD complex. This suggests also that the presence of the NuRD complex at sites occupied by GFI1 reduces transcription relative to sites occupied by GFI1 alone.

### GFI1 affects chromatin remodeling by CHD4 at promoters and enhancers differentially

The active chromatin found here at sites occupied by GFI1 and GFI1/CHD4 complexes in progenitor cells such as GMPs seems at first sight to contradict the established function of both GFI1 and the NuRD complex as transcriptional repressors. In particular, H3K4me2 levels were expected to be depleted at GFI1-occupied sites given its association with LSD1, but this was not always observed when comparing values from aggregation plots. However, it is conceivable that GFI1 or GFI1/NuRD complexes are in a poised state in GMPs and become active only upon differentiation signals that enable them to repress the transcription of genes specific for neutrophil differentiation. Our Metagene analyses, which allowed a quantification and direct comparison between cell types and experiments, provided some clarification and indicated that chromatin openness and H3K4me2 levels at sites occupied by GFI1, CHD4 and GFI1/CHD4 complexes are strongly reduced when GMPs differentiate into preNeu and matNeu cells. In addition, chromatin was more compacted at enhancers bound by CHD4 than at those occupied by GFI1 or GFI1/CHD4. Interestingly, however, H3K4me2 patterns were lowest at the TSS of promoters occupied by GFI1 alone relative to sites occupied by CHD4 or GFI1/CHD4 complexes, which would be in agreement with the presumed function of GFI1 to enable the removal of methyl groups from H3K4 via LSD1. At enhancers, however, H3K4me2 levels are highest at sites occupied by GFI1 compared to sites bound by CHD4 or both GFI1 and CHD4, indicating that the role of GFI1 as a facilitator of chromatin remodeling may be different at promoters and enhancers.

### GFI1, CHD4, and GFI1/CHD4 complexes occupy and regulate different sets of genes

Previous studies have shown that a regulatory network exists between the transcription factors PU.1, IRF8, C/EBPα and GFI1 in myeloid differentiation, and that GFI1 can bind to regions in chromatin together with PU.1 or C/EBPα or IRF8 (Marteijn *et al*, 2007; Olsson *et al*., 2016; Spooner *et al*, 2009). Evidence from a motif analysis suggests that CHD4 affects the role of GFI1 in this regulatory network. While GFI1 and GFI1/C/EBPα complexes bind to sites similarly enriched for IRF and GFI1 consensus binding sites, complexes that contain CHD4 are found at sites enriched for PU.1 binding sites but lacking GFI1 consensus site. This suggests that the presence of CHD4 redirects GFI1 to a different set of promoters, or at least to a different region of a promoter. This also points to the possibility that when in a complex with CHD4, GFI1 may not be required to directly contact DNA or does so through a component of the NuRD complex such as MTA2, which has a DNA binding domain (Denslow & Wade, 2007). The analysis of genes up and down regulated during neutrophil differentiation from GMPs via preNeu to matNeu cells supported this view since it showed that genes occupied by CHD4, GFI1 or GFI1/CHD4 complexes belong to different groups. We propose that in myeloid progenitors GFI1 tethers the NuRD complex through the binding to CHD4 and other components to a specific set of target genes with active or bivalent promoters and active enhancers but remains at a poised state. During neutrophil differentiation, CHD4, GFI1 and GFI1/CHD4 complexes enable the transcriptional regulation of different sets of target genes affecting the immune response, cellular metabolic processes, or nucleosome organization through chromatin remodeling.

## MATERIALS AND METHODS

### BioID-MS data analysis

Peptide searches and protein identification analyses for GFI1_WT-BirA*-Flag or GFI1_ΔSNAG-BirA*-Flag samples were performed as previously described (Bagci *et al*, 2020; Couzens *et al*, 2013; Findlay *et al*, 2018; Shooshtarizadeh *et al*., 2019; Shteynberg *et al*, 2011; Vadnais *et al*., 2018). The BioID-MS data were processed using the ProHits software(Knight *et al*, 2017; Liu *et al*, 2010). The Proteowizard4 tool was used to convert RAW files to .mzXML files. Peptide search and identification were performed by using Human RefSeq version 57 and the iProphet tool integrated in ProHits (Shteynberg *et al*., 2011).

### Dot plot analysis

SAINT files of the wild-type (GFI1WT-BirA*-Flag) and ΔSNAG GFI1 (GFI1ΔSNAG-BirA*- Flag) were processed in ProHits-viz to generate dot plot analyses(Knight *et al*., 2017). Dot plot analyses were performed as described previously(Bagci *et al*., 2020; Shooshtarizadeh *et al*., 2019). Briefly, the Significance Analysis of INTeractome (SAINT) output file of GFI1-BirA*-Flag BioID/MS data generated in ProHits was processed by using the ProHits-viz tools to carry out dot plot analyses(Knight *et al*., 2017; Liu *et al*., 2010) (Choi *et al*, 2011).

### Protein network analysis

Protein network analyses were performed by using Cytoscape(Shannon *et al*, 2003). The SAINT output file of GFI1-BirA*-Flag BioID/MS data was imported to Cytoscape. The existing protein-protein interaction (PPIs) network between preys identified in GFI1-BirA*-Flag BioID/MS screen was imported by using the Biogenet network analysis tool and merged with the GFI1-BirA*-Flag BioID/MS data(Martin *et al*, 2010). The merged protein network was then subjected to MCODE clustering to visualize protein complexes that are connected with the GFI1 bait(Bader & Hogue, 2003).

### Size exclusion chromatography (SEC)

Prior to SEC, 200ul of NE were cleared by centrifugation at 16,000 x g for 10 minutes at 4°C. Cleared NE were size-fractionated on a Superose 6 10/300 column connected to an AKTA-Purifier 10 (Cytiva). Isocratic elution was carried out in 50mM sodium phosphate pH 7, 150mM NaCl, 10% glycerol, 0.5% NP40 and 0.5% Triton-X100, and 500ul fractions were collected. For western blotting analysis of the SEC fractionation, each fraction was precipitated by TCA/DOC, resuspended in 20ul of 2X LDS loading buffer (BioRad), and 80% of each sample was loaded per well on 15 wells 4-15% Mini Protean TGX gels (BioRad) and transferred on PVDF membranes.

### GO term and CORUM analyses

Gene Ontology (GO) and the comprehensive resource of mammalian protein complexes (CORUM) analyses were carried out as described elsewhere using the g:Profiler tool (Reimand *et al*, 2016) (Giurgiu *et al*, 2019; Shooshtarizadeh *et al*., 2019). Briefly, molecular function, biological process or CORUM protein complex analysis of prey interactions recovered in GFI1WT-BirA*-Flag or GFI1ΔSNAG-BirA*-Flag BioID-MS screens are illustrated in heat map analyses. Contaminant proteins such as non-specific interactions or false positives were filtered by using the Contaminant Repository for Affinity Purification (CRAPome) repository prior to the GO term analysis(Mellacheruvu *et al*, 2013). Reviewed UniProtKB entry for each prey protein analyzed in Significance analysis of INTeractome (SAINT) file from ProHits were searched in g:Profiler for GO term or CORUM analysis (Choi *et al*., 2011; Giurgiu *et al*., 2019). GO term enrichment scores were calculated based on the -log10 of corrected *P* values.

### Metagene analysis

The metagene profile of the enrichment of the ATACSeq and H3K4me2 ChIP-Seq experiments in GMP, preNeu and matNeu cell types were produced with previously described bioinformatic procedures(Liang & Keles, 2012). The signal was normalized using the NCIS algorithm(Joly Beauparlant *et al*., 2016).

### Flow cytometry analysis, sorting of GMPs

Hematopoietic cells were analyzed with LSR, or LSR Fortessa flow cytometer (BD Biosciences, Mountain View, CA) and analyzed using BD FACS Diva software (BD Biosciences) or FlowJo (for histogram overlays; Tree Star). For cell sorting, lineage negative BM cells were first depleted using mouse lineage cell depletion kit (Miltenyi Biotec) then applied to five-laser FACSAria II sorter (BD Biosciences).

### Cell culture

K562 (ATCC CRL-3344), HEL (ATCC TIB-180) cells were maintained in RPMI media (Multicell) supplemented with 10% Bovine Growth Serum (RMBIO Fetalgro) and 100 IU Penicillin and 100μg/ml Streptomycin (Multicell). HEK-293 (ATCC CRL-1573) and U2OS (ATCC HTB-96) cells were maintained DMEM media (Multicell) with above-mentioned supplements. We verified that none of the cell lines used in this study were found in the Register of Misidentified Cell Lines maintained by the International Cell Line Authentication Committee

### Western blots

Uncropped and unprocessed scans of the western blots are provided in supplementary files.

### Gene expression profiling by RNA-seq analysis

Bone marrow from 2 tibiae, 2 femora and 2 humeri was harvested in PBS/2.5% FBS and pooled before lineage negative depletion using autoMACS Pro separator (Miltenyi Biotec). Cells were incubated with a lineage antibody cocktail (B220, CD3, CD4, CD8, Gr1, CD11b, NK1.1, Il7R, CD19) and were sorted on FACSAria II sorter (BD Biosciences) to recover GMPs. RNA was extracted using MagMax-96 Total RNA Isolation kit (Ambion) and quality-checked with RNA 6000 Pico kit (Agilent). RNA-seq libraries were prepared from the RNA extracts using the Illumina TruSeq Stranded mRNA Kit according to the manufacturer’s instructions, and sequenced using the TruSeq PE Clusterkit v3-cBot-HS on an Illumina HiSEq 2000 system. Sequencing reads were aligned to the mm10 genome using Tophat v2.0.10(Langmead & Salzberg, 2012). Reads were processed with Samtools(Li *et al*, 2009) and then mapped to Ensembl transcripts using HTSeq(Anders *et al*, 2015). Differential expression was tested using the DESeq R package(Anders & Huber, 2010) (R Core Team 2015, http://www.r-project.org/). A genome coverage file was generated and scaled to RPM using Bedtools(Quinlan & Hall, 2010). RNA-seq data produced for this study are available under accession number GSE173533.

### Functional Analysis

The enrichment of selected biological functions of interest (Supplementary table 1) was also analyzed using the GSEA tool(Subramanian *et al*, 2005). Normalized read counts for Ensembl genes from HTSeq were used and enrichment calculated using 1000 Gene Set permutations. Unsupervised clustering analysis was done using web tool ClustVis (https://biit.cs.ut.ee/clustvis/).

### Consensus Motif Analysis

Motif scanning was performed using the AME tool from the MEME Suite using the JASPAR CORE 2016 database(McLeay & Bailey, 2010).

### Chromatin immuno-precipitation (ChIP)

ChIPs were performed on 1-20x10^6^ sorted cells. The cells were cross-linked with 1.5mM EGS for 20 minutes and 1% formaldehyde for 8 minutes before quenching with 125mM glycine. Cells were lysed in lysis buffer and sonicated using a Covaris E220 to generate 200-600bp fragments(Lee *et al*, 2006). Samples were immuno-precipitated with 2-5 µg of either anti-GFI1 (AF3540, R&D systems), anti-H3K4me1 (ab8895, Abcam), anti-H3K4me2 (ab11946; Abcam), anti-H3K4me3 (ab8580; Abcam) or anti-H3K9me2 (ab1220; Abcam). Libraries were generated according to Illumina’s instructions. Libraries were sequenced on the Illumina Hi-seq 2000 following the manufacturer’s protocols to obtain 50bp paired end reads. External datasets were obtained in the form of .bed files of peaks and .wig visualization tracks, aligned to the mm9 build, except for LSD1, which only included the .bed peak file.

### Annotation databases used

For gene promoters, we used the Ensembl Genes 92 database, dataset GRCh38.p12. (https://useast.ensembl.org/index.html) For enhancer regions, we used the Fantom5 human_permissive_enhancers_phase_1_and_2 enhancers (February 2015) dataset (http://fantom.gsc.riken.jp).

### Statistical Analysis

All p-values were calculated two-sided, and values of p < 0.05 were considered statistically significant. Statistical analysis was done with Graph-Pad Prism software (GraphPad software, La Jolla, CA, USA). The sample size of data points for each assay is shown in Supplementary Data 2.

### Data Availability

The raw proteomics data used in this study are available on ProteomeXchange (http://www.proteomexchange.org) and MassIVE (https://massive.ucsd.edu/ProteoSAFe/static/massive.jsp) under the following accession numbers, respectively: PXD026028 and MSV000087441. The raw ChIP-seq and RNA-seq data, which are presented in Figures 4 and Supplementary Figures 4, have been uploaded to the GEO Datasets repository (https://www.ncbi.nlm.nih.gov/gds) and is available under the following accession numbers: GSE173533. Previously published ChIP-seq and ATAC-Seq data, which is presented in Figure 4 and Supplementary Figure 4, are available under the following accession numbers: H3K27ac (GSM1441273), H3K27me3 (DRR023959), H3K9me3 (DRR023962) and ATAC-Seq (DRR023962).

## ACKNOWLEDGEMENTS

We are indebted to Mathieu Lapointe for technical assistance, Marie-Claude Lavallée and Jo-Anny Bisson for excellent animal care, Eric Massicotte and Julie Lord for FACS and cell sorting. We thank the Genome Quebec for performing HT sequencing and Anne Helness for ChIP-Seq and ATAC-Seq experiments. Tarik Möröy holds a Canada Research Chair (Tier 1) and grants from the CIHR (MOP-84238, MOP-94846, FDN-148372) and the Cancer Research Society. Charles Vadnais was supported by fellowships from the Fonds de recherche Quebec – Santé (FRQS) and from the CIHR. Halil Bagci was supported by a doctoral training award from FRQS (#33603) and NSERC Discovery Grant (RGPIN-2016-04808 to Jean-François Coté). Jean-François Coté holds the TRANSAT chair in breast cancer research. Christian Trahan was supported by CIHR Project grant (PJT-153313 to Marlene Oeffinger).

## AUTHORS AND CONTRIBUTIONS

Anne Helness: Concept and design, Collection of data, data analysis, manuscript writing

Jennifer Fraszczak: Collection of data, data analysis, manuscript writing

Christian Trahan: Collection of data, data analysis, manuscript writing

Charles Joly-Beauparlant: Analysis of genomic data

Peiman Shooshtarizadeh: Concept and design, Collection of data, data analysis

Marina Ayoub: Collection of data, data analysis

Kaifee Arman: Collection of data, data analysis

Halil Bagci: Proteomic data analysis, manuscript writing

Jean-François Coté: Concept and manuscript writing, reagents

Arnaud Droit: Concept and analysis of genomic data

Marlene Oeffinger: Concept and manuscript writing

Tarik Möröy: Concept and design, data analysis and supervision, manuscript writing, final approval, provision of funds

## Supplementary Material

**Extended View Figure 1:**

**A)** Amino acid sequences of CHD4 and MTA2 with peptides identified through mass spectrometry highlighted in yellow with the corresponding spectra below.
**B)** Immunoprecipitation experiments with cells expressing Flag tagged GFI1 to detect MTA2 in the presence of absence of Benzonase
**C) -D)** Heatmaps showing CORUM terms, Biological processes and Molecular functions of associated protein complexes, respectively, of related GO terms in Flp-In T-REx HEK293 cells expressing GFI1WT-BirA*-Flag or GFI1ΔSNAG-BirA*-Flag. GO term or CORUM protein complex enrichment scores are shown as the -log10 of corrected P values, illustrated by different color intensities.

**Extended View Figure 2:**

**A)** Schematic depiction of the *Csf1* locus encoding M-CSF. Shown is the enrichment of reads after ChIP-seq with antibodies against GFI1 or CHD4 in GMPs from either WT or GFI1KO mice and the enrichment of reads after an RNA-seq experiment from WT of GFI1 KO GMPs. The transcription start site is indicated (TSS).
**B)** ChIP-qPCR for CHD4 in primary GFI1 WT and KO GMPs of seven exemplary genes selected from the group of 841 loci where CHD4 binding was lost in the absence of GFI1.
**C)** RNA-seq profile of the *Chd4* gene in GMPs from WT or *Gfi1* KO mice
**D)** Flow cytometric profile for CHD4 (intracellular staining) in GMPs from WT or *Gfi1* KO mice

**Extended View Figure 3**

Compilation of RNA-seq and ChIP seq data for known GFI1 target genes(Horman *et al*, 2009) that are either upregulated (A) or downregulated (B) in *Gfi1* KO GMPs

**Extended View Figure 4**

Depiction of the ATAC-Seq and ChIP-Seq data at the GFI1 target gene *Csf1*. Indicated are the tracks corresponding to the individual experiments, the gene and the TSS.

**Extended View Figure 5**

**Gfi1:GFP expression in cellular subsets of the neutrophil lineage**

**A)** Flow cytometric sorting strategy for pre-neutrophils (preNeu), immature Neutrophils (immatNeu) and mature Neutrophils (matNeus) cells according to the following markers: preNeu: Mac1^+^Gr1^+^ckit^+^CXCR4^+^, immNeu: Mac1^+^Gr1^+^CXCR4^-^cKit^-^SiglecF^-^CXCR2^-^, matNeu: Mac1^+^Gr1^+^CXCR4^-^cKit^-^SiglecF^-^CXCR2^+ (^Evrard *et al*., 201^8^).
**B)** Western blot for GFI1 and Lamin expression using nuclear extracts from the cell populations indicated in **A)**
**C)** GFP intensity of the neutrophil lineage cell sub populations of total bone marrow from heterozygous Gfi1:GFP knockin mice, expressing a GFP cDNA under the control of the Gfi1 promoter (Yucel *et al*., 2004). GMP: lin^-^ckit^+^Sca1^-^CD16/32^+^CD34^+,^ Pre-neutrophils: Gr1^+^Mac1^+^Ckit^+^CXCR4^+,^ immature Neutrophils: Gr1^+^Mac1^+^ckit^-^CXCR4^-^SiglecF^-^ CXCR2^-^, Mature Neutrophils: Gr1^+^Mac1^+^ckit^-^CXCR4^-^SiglecF^-^CXCR2^+^, MEP (lin^-^ Ckit^+^Sca1^-^CD16/32^-^CD34^+^), which do not express GFI1(Zeng *et al*., 2004), are used as a control.

**Extended View Fig. 6**

**A) B)** RNA-seq normalized reads of genes associated with chemotaxis or phagocytosis according to(Evrard *et al*., 2018)

**Extended View Fig. 7**

**A) - C)** RNA-seq normalized reads of genes associated with myeloid differentiation according to(Evrard *et al*., 2018)

**Supplementary Table 1.** Unfiltered SAINT result of the GFI1_WT, GFI1_DeltaSNAG and GFI1B BioID data

**Supplementary Table 2.** Biological processes, Molecualr Functions and CORUM protein complexes

**Supplementary Table 3.** P-values and FDR values for Metagene analysis shown in Figure 6

| group1 | group2 | pvalue | fdr |  |
| --- | --- | --- | --- | --- |
| H3K4me2_GMP_CHD4 | H3K4me2_GMP_CHD4/Gfi1 | 0,388959964 | 0,400073106 | Promoter |
| H3K4me2_GMP_CHD4 | H3K4me2_GMP_Gfi1 | 3,0832E-05 | 9,2496E-05 |  |
| H3K4me2_GMP_CHD4/Gfi1 | H3K4me2_GMP_Gfi1 | 0,00268003 | 0,004824053 |  |
| H3K4me2_PreNeu_CHD4 | H3K4me2_PreNeu_CHD4/Gfi1 | 0,024944722 | 0,029933666 |  |
| H3K4me2_PreNeu_CHD4 | H3K4me2_PreNeu_Gfi1 | 1,98228E-09 | 1,01946E-08 |  |
| H3K4me2_PreNeu_CHD4/Gfi1 | H3K4me2_PreNeu_Gfi1 | 2,88873E-07 | 1,15549E-06 |  |
| H3K4me2_MatNeu_CHD4 | H3K4me2_MatNeu_CHD4/Gfi1 | 0,00754659 | 0,012348966 |  |
| H3K4me2_MatNeu_CHD4 | H3K4me2_MatNeu_Gfi1 | 0,019114389 | 0,026466077 |  |
| H3K4me2_MatNeu_CHD4/Gfi1 | H3K4me2_MatNeu_Gfi1 | 0,000110681 | 0,000267533 |  |
| ATACSeq_GMP_CHD4 | ATACSeq_GMP_CHD4/Gfi1 | 5,45645E-05 | 0,000151102 |  |
| ATACSeq_GMP_CHD4 | ATACSeq_GMP_Gfi1 | 1,16249E-32 | 1,39499E-31 |  |
| ATACSeq_GMP_CHD4/Gfi1 | ATACSeq_GMP_Gfi1 | 0,021706006 | 0,028382655 |  |
| ATACSeq_PreNeu_CHD4 | ATACSeq_PreNeu_CHD4/Gfi1 | 0,000111472 | 0,000267533 |  |
| ATACSeq_PreNeu_CHD4 | ATACSeq_PreNeu_Gfi1 | 1,83812E-43 | 3,30862E-42 |  |
| ATACSeq_PreNeu_CHD4/Gfi1 | ATACSeq_PreNeu_Gfi1 | 0,00030706 | 0,000650244 |  |
| ATACSeq_MatNeu_CHD4 | ATACSeq_MatNeu_CHD4/Gfi1 | 0,000971985 | 0,00194397 |  |
| ATACSeq_MatNeu_CHD4 | ATACSeq_MatNeu_Gfi1 | 1,01358E-45 | 3,64888E-44 |  |
| ATACSeq_MatNeu_CHD4/Gfi1 | ATACSeq_MatNeu_Gfi1 | 2,95617E-06 | 9,67472E-06 |  |
| H3K4me2_GMP_CHD4 | H3K4me2_GMP_CHD4/Gfi1 | 0,022075398 | 0,028382655 | Enhancer |
| H3K4me2_GMP_CHD4 | H3K4me2_GMP_Gfi1 | 1,1483E-17 | 6,88981E-17 |  |
| H3K4me2_GMP_CHD4/Gfi1 | H3K4me2_GMP_Gfi1 | 0,000274254 | 0,000617071 |  |
| H3K4me2_PreNeu_CHD4 | H3K4me2_PreNeu_CHD4/Gfi1 | 0,001469442 | 0,002784206 |  |
| H3K4me2_PreNeu_CHD4 | H3K4me2_PreNeu_Gfi1 | 5,39382E-28 | 3,88355E-27 |  |
| H3K4me2_PreNeu_CHD4/Gfi1 | H3K4me2_PreNeu_Gfi1 | 1,60435E-06 | 5,77566E-06 |  |
| H3K4me2_MatNeu_CHD4 | H3K4me2_MatNeu_CHD4/Gfi1 | 0,005913535 | 0,010137489 |  |
| H3K4me2_MatNeu_CHD4 | H3K4me2_MatNeu_Gfi1 | 2,18322E-28 | 1,9649E-27 |  |
| H3K4me2_MatNeu_CHD4/Gfi1 | H3K4me2_MatNeu_Gfi1 | 1,10396E-08 | 4,96783E-08 |  |
| ATACSeq_GMP_CHD4 | ATACSeq_GMP_CHD4/Gfi1 | 0,012029081 | 0,018828127 |  |
| ATACSeq_GMP_CHD4 | ATACSeq_GMP_Gfi1 | 0,408054971 | 0,408054971 |  |
| ATACSeq_GMP_CHD4/Gfi1 | ATACSeq_GMP_Gfi1 | 0,091715442 | 0,103179872 |  |
| ATACSeq_PreNeu_CHD4 | ATACSeq_PreNeu_CHD4/Gfi1 | 0,013266204 | 0,019513086 |  |
| ATACSeq_PreNeu_CHD4 | ATACSeq_PreNeu_Gfi1 | 0,024441512 | 0,029933666 |  |
| ATACSeq_PreNeu_CHD4/Gfi1 | ATACSeq_PreNeu_Gfi1 | 0,369143642 | 0,390857973 |  |
| ATACSeq_MatNeu_CHD4 | ATACSeq_MatNeu_CHD4/Gfi1 | 0,013550754 | 0,019513086 |  |
| ATACSeq_MatNeu_CHD4 | ATACSeq_MatNeu_Gfi1 | 0,077456047 | 0,089948958 |  |
| ATACSeq_MatNeu_CHD4/Gfi1 | ATACSeq_MatNeu_Gfi1 | 0,313178777 | 0,341649574 |  |

